# Neural dynamics of an extended frontal lobe network in goal-subgoal problem solving

**DOI:** 10.1101/2025.05.06.652442

**Authors:** Valentina Mione, Jascha Achterberg, Daniel J. Mitchell, Makoto Kusunoki, Mark J. Buckley, John Duncan

## Abstract

In goal-directed behavior, cognitive control calls for representation of current state, goal state, and current move, often with hierarchical organization into a series of steps or subgoals. In the human brain, a network of “multiple-demand” or MD regions underpins cognitive control, and core task features are likely encoded and integrated across this distributed network. Here we recorded from four putative homologs to human MD regions in the frontal cortex – ventrolateral (vlPFC), dorsomedial (dmPFC), dorsal premotor (dPM) and anterior insula/orbitofrontal (I/O) cortex - as two male monkeys solved a multi-step, on-screen spatial maze. Across regions there was substantial overlap but also wide quantitative variation in encoding task variables. Sensory input and current state were strongly coded in vlPFC, goal most stably in dmPFC, and move most rapidly in vlPFC and dPM. I/O responded during revision of a prepared route. In all regions, an abstract code of problem structure marked progress from problem start to end, with hierarchical separation of progress within and between successive steps. Partial specialisations, we suggest, arise from distinct connectivity, while broad co-recruitment reflects strong information exchange. Our results show how core components of goal-directed behavior are encoded across a distributed network of partial frontal lobe specialists, putatively homologous to frontal components of the human MD system.

## Introduction

It is widely accepted that the frontal lobes are involved in high-level cognitive control. Control must rest on an internal model of a task’s contents and requirements ^1^ ^2^. This model must include a specification of current state, goal state, and actions chosen to reduce the difference between the two ^3^ ^4^. Often behavior has a hierarchical structure, with a higher goal decomposed into a sequence of lower-level subgoals ^3^ ^4^. Preparing breakfast, for example, may involve subgoals such as making coffee, toasting bread, and buttering the toast. All these aspects of the internal model must be integrated to direct behavioral choice. The ability to create very many complex wholes from a sequence of simpler parts underlies the compositionality and productivity of cognition.

In human brain imaging, cognitive control is robustly linked to a cortical/subcortical “multiple demand” (MD) network, recruited during many kinds of cognitive activity ^5^ ^6^. Core components of this network are now tightly defined in several regions of lateral and dorsomedial frontal cortex, in the anterior insula, and within the intraparietal sulcus ^7^, as well as regions of caudate, thalamus and cerebellum. Brain imaging data show frequent co-recruitment, strong functional connectivity, yet quantitative variations in activity pattern across the components of this network ^7^. Widespread components of this network likely bring local access to information of many kinds. At the same time, strong functional connectivity suggest integration, combining the components of a cognitive operation ^7^ ^8^ ^6^.

In nonhuman primates, many studies document coding of diverse task features in lateral prefrontal, dorsomedial prefrontal, dorsal premotor and insular cortex ^9^ ^10^ ^11^ ^12^ ^13^ ^14^ ^15^ ^16^ ^17^ ^18^ ^19^ ^20^. Between regions, neural properties often overlap, though with quantitative differences (e.g. ^9^ ^11^ ^12^ ^13^ ^17^ ^18^). Differences likely reflect different anatomical connections ^21^, while dense connectivity within the frontal lobe suggests a medium for extensive information exchange ^22^ ^23^. The findings suggest that cognitive control involves a distributed, interactive ^24^ network of multiple frontal lobe regions. Across this network, however, we still lack an integrated overview of shared and specialised functions encoding current state, goal state, move and hierarchy.

To investigate this question, we trained two macaques to solve problems in an on-screen maze. Each problem had a hierarchical structure, with a flexible series of subgoals (intermediate maze locations) selected on the path to the final goal. To separate coding of higher goal and component steps, different routes could lead to the same final goal, and different goals could be reached by partially overlapping routes. We recorded from four separate regions of frontal cortex, aiming for tentative homologs to human MD regions. Across our extended frontal network, we examined the dynamics of specialised and overlapping neural coding for current state, goal, chosen moves and abstract problem structure.

## Results

The task is illustrated in Figure 1A. In each problem, the monkey began by fixating the center location in a 5 x 5 grid of lines, presented on a computer screen (Figure 1a, left, green dot). Intersections between these lines defined 5 x 5 permitted fixation locations as each problem was solved. With a series of saccades, the task was to shift gaze to one of four goal locations (Figure 1A, blue dots). Each saccade moved just one step through the maze, i.e. from the currently fixated location to an immediately adjacent location (intersection). By controlling available choices at each step, we created a total of 16 possible routes through the maze, 8 reaching the goal in just 2 steps (Figure 1a, orange lines), and 8 requiring 4 steps (Figure 1A, green lines).

**Figure 1.**
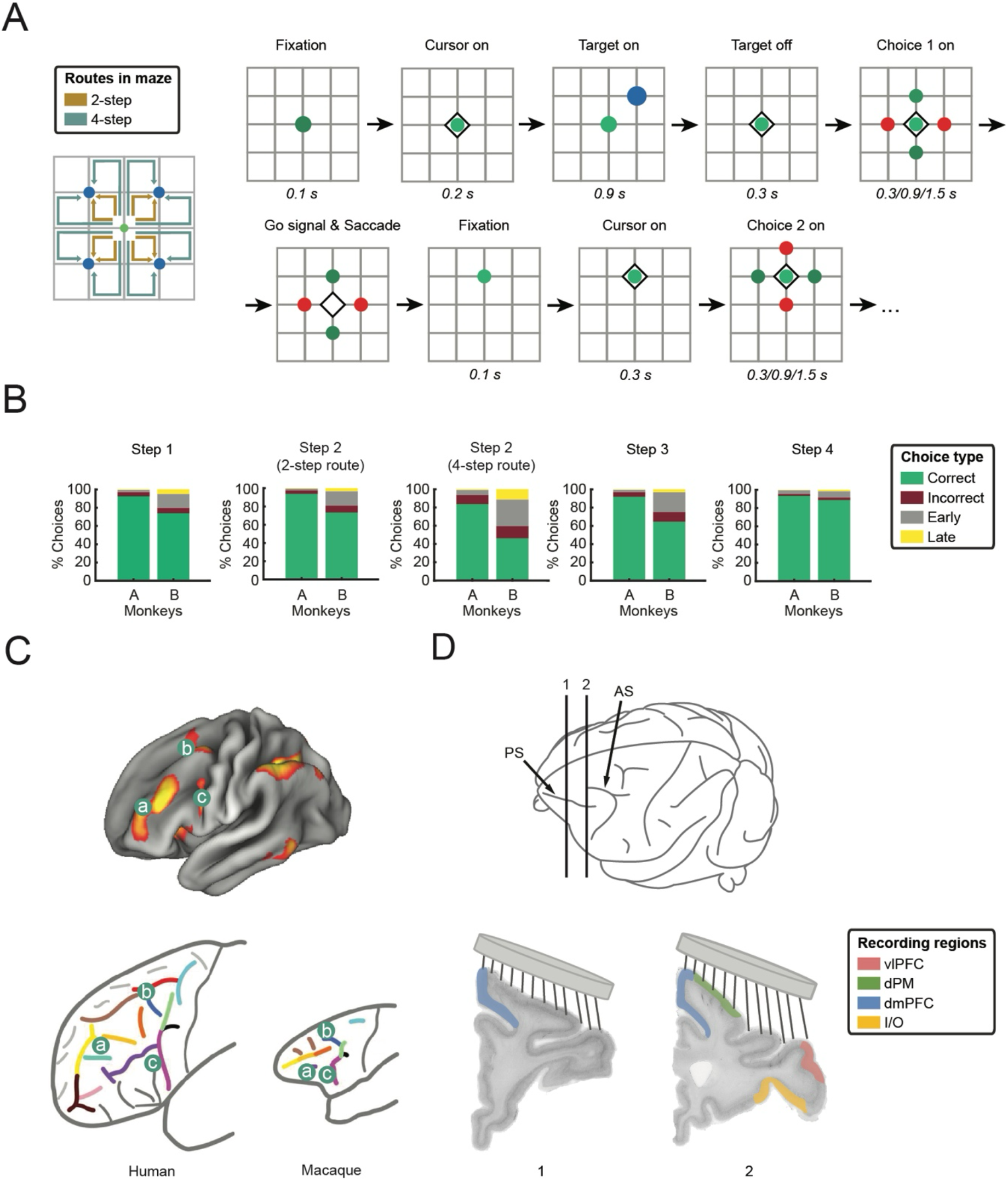
A. Task. Left: The maze was presented as a grid of lines on a computer screen. The 5 x 5 maze locations were defined by intersections between these lines. At each step, the monkey made a saccade from current location (fixation point) to an immediately adjacent location. Green dot - starting location; blue dots – possible goal locations; arrows – 16 possible routes from start to goal (orange - 2-step; green - 4-step). Right: Timing of events from problem start to step 2 choice stimulus. Similar timings were used for each remaining step. Diamond and dots larger than actual size on maze. B. Behavior. For each step, data are based on problems for which the corresponding choice stimulus was presented, i.e. excluding problems aborted at an earlier point. C. Correspondence analysis, suggesting putative match between human MD regions on lateral frontal surface and corresponding regions in macaque brain. Above – MD activity from ^7^, projected on partially inflated cortical surface. Green dots with letters mark major MD regions. Below – putative sulcal correspondences between human and macaque, copied from ^25^. D. Recording regions illustrated on example coronal brain slices. Disc above each slice shows approximate extent of recording chamber, with schematic electrodes indicating approximate recorded area. PS – principal sulcus; AS – arcuate sulcus.

Timing details are shown in Figure 1A (right). At the start of each problem, once central fixation was acquired, a cursor (diamond) joined the central green fixation point. This was followed by a cue indicating the goal for the current problem (Figure 1A, left, one of the four locations marked by blue dots). The goal cue was shown for 900 ms, followed by a delay and then a pair of alternative moves for the first step. Of the four locations (grid intersections) adjacent to the start location, two (varying from problem to problem) were coloured green, indicating that they were available for selection, while the other two were coloured red, indicating unavailability. Following a further delay, the monkey received a go signal (disappearance of central fixation point), and could made a saccade to one of the two green alternatives, the correct choice being the one moving closer to the goal. This process was then repeated for step 2: green dot and cursor marking currently fixated location, delay, presentation of available and unavailable choices, further delay, go signal and saccade. For 2-step problems, one of the step 2 alternatives was the final goal, allowing the monkey to saccade to this goal for smoothie reward (Figure 1A, left, orange arrows). For 4-step problems, in contrast, the goal position was unavailable (red), and the monkey was forced to make a step to a green choice on the outer perimeter of the maze. Two further choices, with similar timing, were required before the final goal was reached (Figure 1A, left, green arrows). The problem was always aborted for any error (see Methods), meaning that reducing numbers of trials were completed for each successive step.

Behavioral results for the two animals are shown in Figure 1B. Except that monkey B often failed to wait for the go signal, the performance of both animals was accurate at each step. Discarding early and late responses, averaging across steps and problem types, the mean probability of correct choice was 0.95 for monkey A, 0.89 for monkey B.

Using a semi-chronic recording system, we recorded neural activity over 72 sessions in monkey A, 70 sessions in monkey B, with different groups of neurons recorded in each session. To target recordings we used current knowledge of human MD regions, combined with a recent analysis of correspondences between the frontal lobes of human, chimpanzee, baboon and macaque ^25^. On the lateral frontal surface, MD activity extends forward from the inferior frontal junction (Figure 1C, upper; regions a and c) ^7^. The correspondence analysis (Figure 1C, lower) places the inferior frontal junction within the macaque ventrolateral prefrontal cortex (vlPFC; Figure 1C; region c), and based on a suggested correspondence between the macaque principal sulcus and the human posteromedial frontal sulcus, we tentatively match more anterior human MD regions also to macaque vlPFC (Figure 1C; region a). A further human MD patch is posterior and dorsal on the lateral frontal surface (Figure 1C, upper; region b), and the correspondence analysis suggests match to a region of dorsal premotor cortex (dPM), behind the upper limb of the arcuate sulcus. On the dorsomedial surface, the human MD region overlaps presupplementary motor area (pre-SMA) and dorsal anterior cingulate; as a putative match we selected a region of dorsomedial prefrontal cortex (dmPFC) centered on the macaque pre-SMA. Finally, a highly consistent human MD region is found in the anterior insula and adjacent frontal operculum; as a tentative match, we recorded from a region of intermediate agranular insula and immediately adjacent orbitofrontal cortex (I/O). Based on the atlas of Saleem and Logothetis ^26^, our selected recording regions (Figure 1D) were thus vlPFC, including the anterior bank of the ventral arcuate sulcus (315 neurons, 236/79 from monkeys A/B, regions 45, 44, 8A-F5), dPM (341 neurons, 318/23 from monkeys A/B, regions F2, F7), dmPFC (360 neurons, 311/49 from monkeys A/B, regions F3, F6, 9m/8Bm), and I/O (358 neurons, 327/31 from monkeys A/B, regions Iai, 12o, 13l). For main analyses below, data from the two animals were pooled. For vlPFC and dmPFC only, given adequate neural samples from both animals, we show broad similarity in data from the two.

The following sections describe three kinds of analyses. One important aspect of a task model is provided by sensory input, indicating current environmental state and its constraints on behavioral options. In the first analysis, we compared regions for their response to three kinds of sensory input. In the second analysis, we took advantage of our task’s separation of final goals from component moves. Using principal component analysis (PCA), we defined separate representational spaces for goal and move, and examined the dynamics of representation in each space. In the third analysis, we examined how patterns of activity evolve from problem start to end, reflecting an abstract code of task structure, independent of individual goals and moves.

### Sensory input

For a first comparison of regions, we examined responses to sensory input. In our maze problems, unpredictable input occurred at three points: at initial presentation of the goal, at step 1 when available alternatives indicated a horizontal or vertical first move, and at step 2 when alternatives indicated a 2- or 4-step route. In each case, for each individual neuron, we used analysis of variance (ANOVA) on spike rate to measure proportion of explained variance (PEV) for goal location during goal presentation, for horizontal/vertical choice available at step 1, and for 2- vs 4-step at step 2 (see Methods). ANOVAs were conducted on data from successive 50 ms windows, beginning 200 ms before each stimulus event. Randomisation tests (see Methods) were used to identify significant periods of information coding (PEV > 0) in each region, and to compare PEV between regions. Here and subsequently, analyses for each step included only data from trials on which this step was completed correctly. Mean values of PEV across all recorded neurons in each region are shown in Figure 2. For goal (Figure 2, left), an early peak in vlPFC, first significant in the window 50-100 ms post onset and maximal in the period 100-200 ms, was followed by rapid decline to a plateau lasting to the end of the goal presentation. In other regions, the peak developed more slowly, and most weakly in I/O, but again, goal coding remained largely significant to the end of the presentation period. For horizontal/vertical first move (Figure 2, middle), vlPFC again showed the strongest early coding, now beginning in the window 150-200 ms post onset of the choice alternatives, but by 250-300 ms, there was substantial convergence between vlPFC, dPM and dmPFC. For 2- vs 4-step coding at step 2 (Figure 2, right), coding onset was again fastest in vlPFC (150-200 ms window), but thereafter, coding became strongest in dmPFC and, slightly later, in I/O.

**Figure 2.**
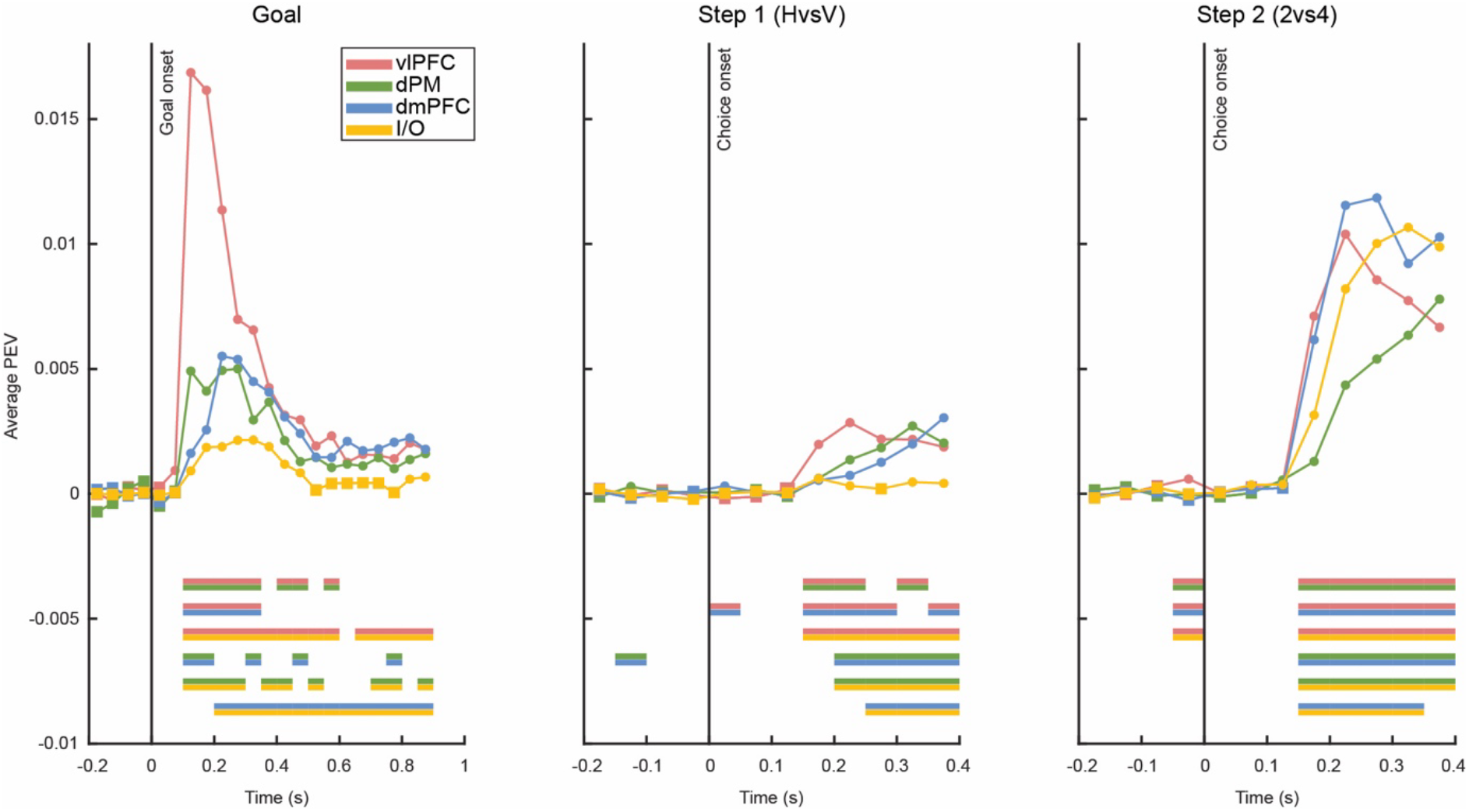
Mean proportion of explained variance (PEV) for goal (left, aligned at goal onset), horizontal (H) versus vertical (V) first move (middle, aligned at onset of choice 1 stimulus), and 2-versus 4-step problem (right, aligned at onset of choice 2 stimulus). For each region, values of PEV significantly > 0 (*p* < .05, FDR corrected) are indicated by round dots, nonsignificant by square dots. Horizontal bars (bottom) show significant differences between each pair of regions.

These three types of sensory input called for different kinds of computation. In line with human fMRI data, showing coactivation of MD regions, the results show substantial overlap in response of the four regions. At the same time they show clear functional separations, with vlPFC showing strong and rapid responses to all novel sensory input, and I/O strongly responsive only to an input differentiating immediate goal access from a longer, more demanding path.

For vlPFC and dmPFC, separate data for the two animals are shown in Supplementary Figure 1. Both animals showed early vlPFC responses to all three types of sensory input, though for monkey B, step 1 horizontal/vertical discrimination, the early difference between regions was not significant.

### Location spaces

For the next analyses we wished to define separate representational spaces for two aspects of the maze problem. Specifically, we aimed to separate a representation of the problem goal – required to direct a series of component choices - from representation of individual movements.

We used combined data from steps 2 (2-step problems only) and 4, since combining these two steps, each possible goal (top left, top right, bottom left, bottom right) is associated with each possible move/saccade (left, right, up, down). For each goal, over a window extending from -200 to +100 ms from choice stimulus presentation, we averaged firing rates for each neuron over the four possible moves. ANOVAs showed neurons with significant (*p* < .05) goal discrimination in all four frontal regions (27%, 31%, 21% and 16% respectively in vlPFC, dPM, dmPFC, I/O). For monkeys A/B, values for vlPFC were 31%/16%, and for dmPFC 21%/20%.

For each region, using combined data from the two animals, we then used PCA to extract a representational space differentiating the four goal locations. Note that different goal locations were partially associated with different current locations (at steps 2 and 4, current location always immediately adjacent to final goal). Movements in the representational space could accordingly be driven by either goal location or current location. For each region, we used data from steps 1 to 4 to indicate the relative prominence of these two.

For each region, reflecting the two-dimensional layout of the maze, the first two PCs accounted for the great majority of variance (89, 91, 90, 84% respectively for vlPFC, dPM, dmPFC and I/O). For each region, for the combined data from steps 2 and 4, projections onto the first two PCs showed the expected structure, with the four locations organized in a layout approximately matching their spatial layout on the screen (Figure 3, top panel for each region, “master projection”).

With this space defined, we proceeded to examine projections from other problem periods. For each of the four steps separately, we took data from three periods: -200 to +100 ms from choice stimulus presentation (pre-choice), as used in the original creation of the spaces; +200 to +400 ms from choice stimulus (choice), to capture the movement selection period; and -200 to +100 ms from the go signal (go). For each region, the resulting projections for each goal are shown in the lower four rows of Figure 3, with the step 2 row using only data from 2-step trials (for similar results in 4-step trials, see Supplementary Figure 2). In each row, the rightmost plot shows cross-validated proportion of variance explained by projections into the PCA space (see Methods).

For vlPFC (Figure 3, top left), there was a striking and unique pattern. At step 1, when the monkey always fixated the center location, projections of the four goal locations were not significantly different. Significant differences appeared from steps 2 to 4, especially in the pre-choice period of each step, but were by far the strongest at step 4. Notably, at step 4, current locations were maximally different for the four goals (outer ring locations adjacent to each goal). This pattern is suggestive of a strong code for current location in the maze, available to the animal through current sensory input.

For dmPFC (Figure 3, bottom left) there was a complementary pattern. For this region, the four goals were significantly and consistently separated from the pre-choice period of step 1 to the go period of step 4. A corresponding analysis of data from the early target presentation period (100 to 400 ms post onset of the goal stimulus at trial start) showed strong separation even here (proportion of explained variance 0.30), much greater than for other regions (0.05 to 0.08). For dmPFC, these results suggest a consistent representational space separating the four goals, established immediately at target presentation and maintained to problem end.

**Figure 3.**
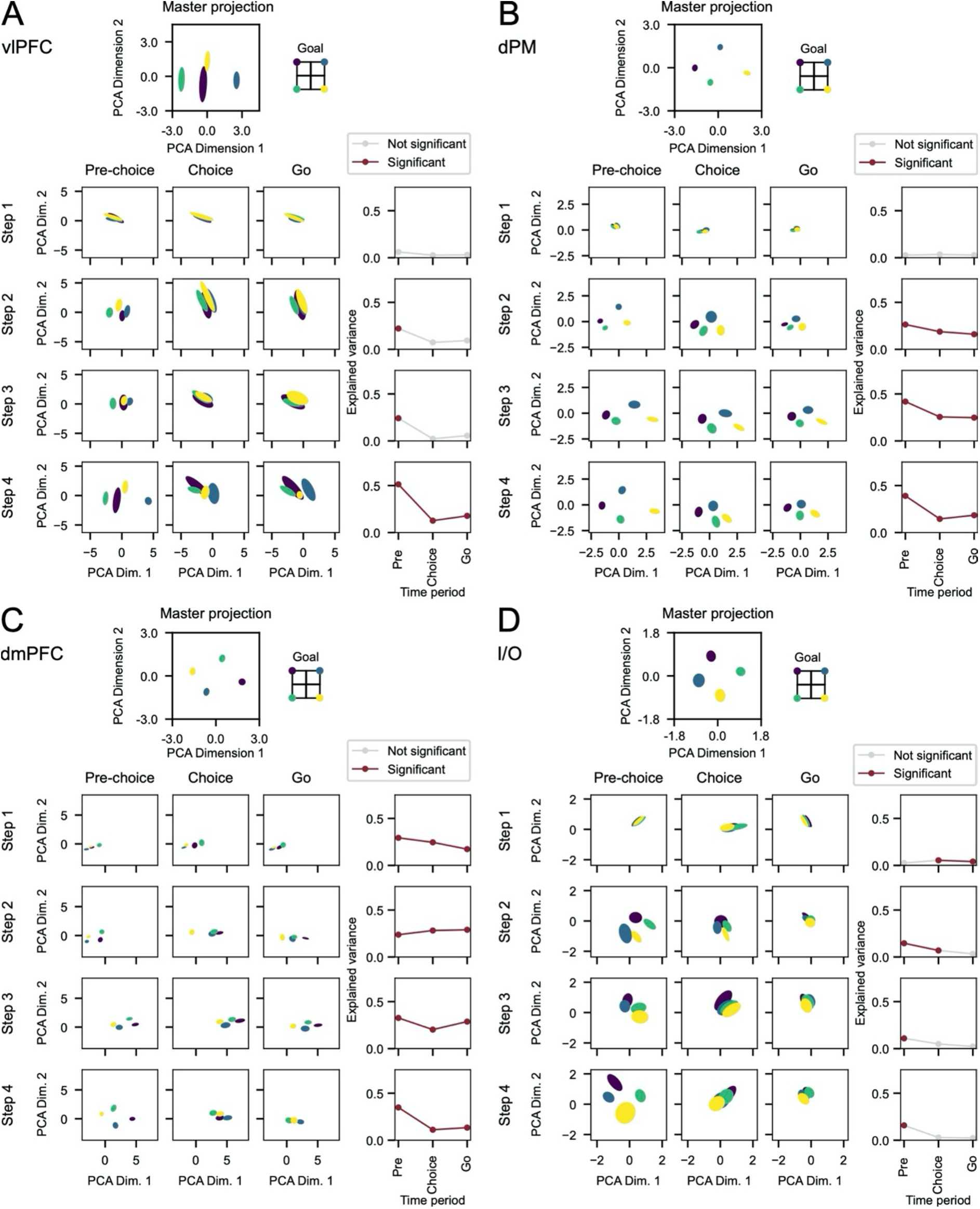
Location spaces for A - vlPFC, B - dPM, C – dmPFC, D – I/O. For each region, the top panel shows projections into a master space derived from data averaged over four moves for each goal location (pre-choice periods from steps 2 (2-step problems only) and 4). Lower rows show projections into this master space for data from individual time windows in each step (left three panels), with proportion of explained variance in each window to the right (purple, *p* < .05, FDR corrected). Step 2 row takes data only from 2-step problems. Ellipses show confidence intervals derived by bootstrap resampling.

For dPM and I/O, results were a mixture of these two patterns. Again, projections of the four goals were closely similar at step 1, but then separated across later steps, again most strongly in the pre-choice period.

To further differentiate representations of goal and current location, we used data from the pre-choice period of steps 2 (2-step problems) and 3 (4-step problems). For these periods, each goal was associated with one of two current locations, and each current location was associated with one of two goals. Projections of the resulting eight cases are shown in Figure 4, separately classified by current location and goal. Beneath these projections are shown mean distances between points with same goal/different current location, same current location/different goal, and as a comparison, both different (baseline). (Note that, for the first two cases, required moves were always different; for comparability, we also selected baseline pairs with different moves.) For vlPFC, distances were shortest for same current location/different goal, supporting the conclusion that, in this region, the coding space was predominantly concerned with current location. For dmPFC there was a complementary result, with shortest distance for same goal/different current location. For the remaining two regions, results suggested representation of both current location and goal.

Results for vlPFC are intriguing because, though goal location was strongly represented during initial target presentation (Figure 2), this did not translate into a later, stable goal space. This conclusion was confirmed by constructing a new goal space from the target presentation period (+100 to +400 ms from goal onset), and examining how much between-goal variance this space could explain in pre-choice periods from each subsequent step. Proportion explained variance was always below 0.13, and nonsignificant except for the period preceding step 2. To a large extent, in vlPFC, the strong goal representation created during target presentation was lost in subsequent steps towards this goal.

**Figure 4.**
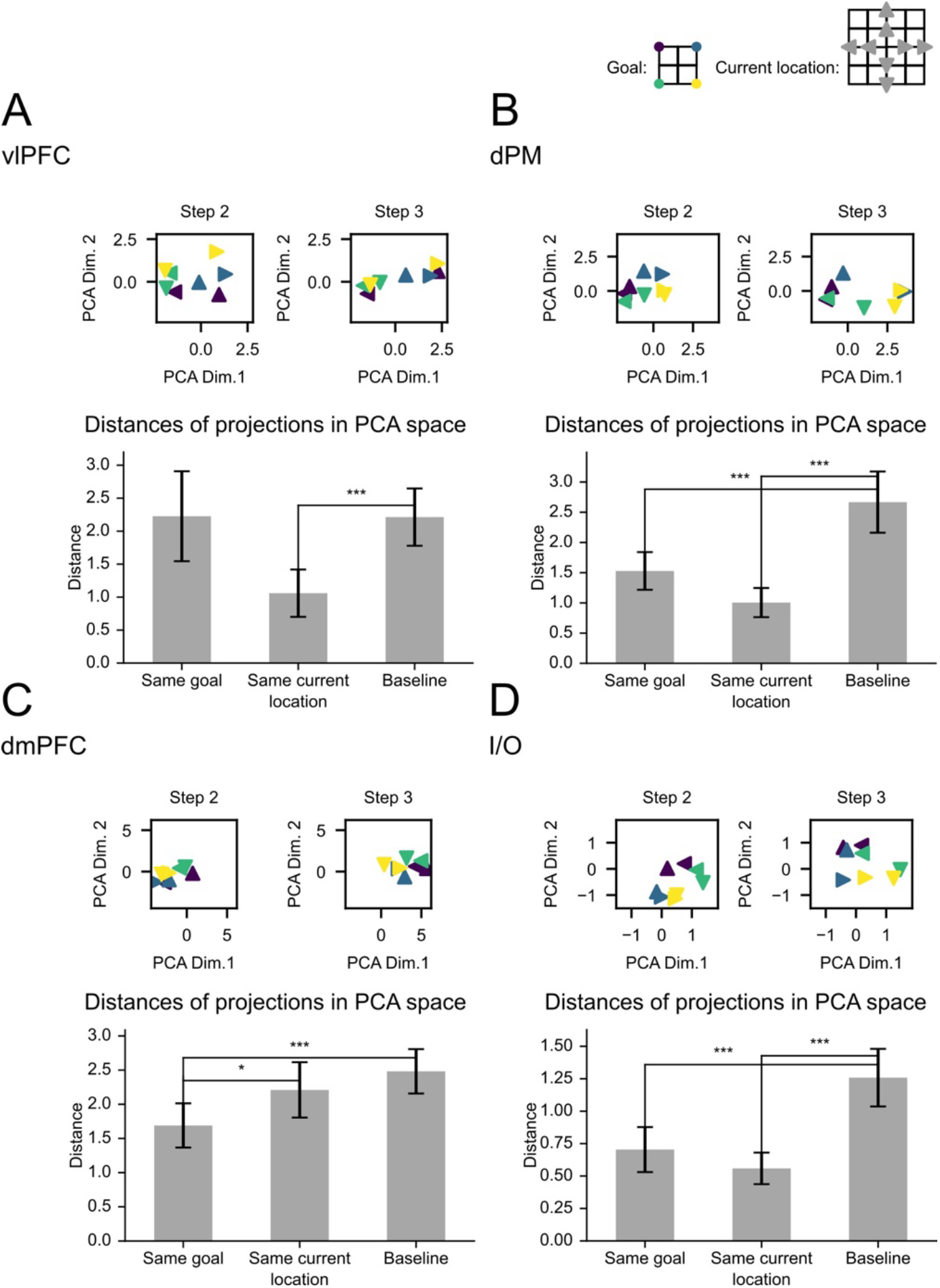
Projections into master location space for pre-choice data from steps 2 (2-step problems) and 3. A - vlPFC, B - dPM, C – dmPFC, D – I/O. For each region, top panels show projections for data sorted by goal location x current location. Bottom shows mean distance between projections for pairs of points with same goal (different current location and move), same current location (different goal and move), and baseline (different goal, current location and move). Whiskers show +/- 1 standard deviation over bootstrap samples. * *p* < .05; *** *p* < .001.

Together, these results show a mixture of shared and specialised coding. Most striking was differential coding in vlPFC and dmPFC. dmPFC was unique in carrying a stable goal code from problem start to end. The stability of this goal code corresponds to the stable role of the final goal in directing a series of component moves. Strong coding of current location in the vlPFC again suggests a dominant influence of immediate visual input. In dPM and I/O, there was evidence for codes of both current and goal locations.

### Move spaces

To define a move space we took a complementary approach. Again we combined data from steps 2 (2-step problems only) and 4, so that for each move, we could average data across the four different goals. To appropriately capture move selection we used the post-choice period, +200 to +400 ms from choice stimulus onset. ANOVAs showed neurons with significant (*p* < .05) move discrimination in all four frontal regions (48%, 37%, 26% and 17% respectively in vlPFC, dPM, dmPFC, I/O). For monkeys A/B, values for vlPFC were 55%/29%, and for dmPFC 27%/20%.

Again, PCA used combined data from the two animals. For each region, once again, the great majority of variance was captured by the first two principal components (86, 88, 91, 76% respectively for vlPFC, dPM, dmPFC and I/O), and projections for the four possible moves approximated their objective spatial relations (Figure 5, top part of each panel, “master projection”). Confidence intervals (Figure 5, ellipses; see Methods) showed that move directions were discriminated much more weakly in I/O than other regions.

**Figure 5.**
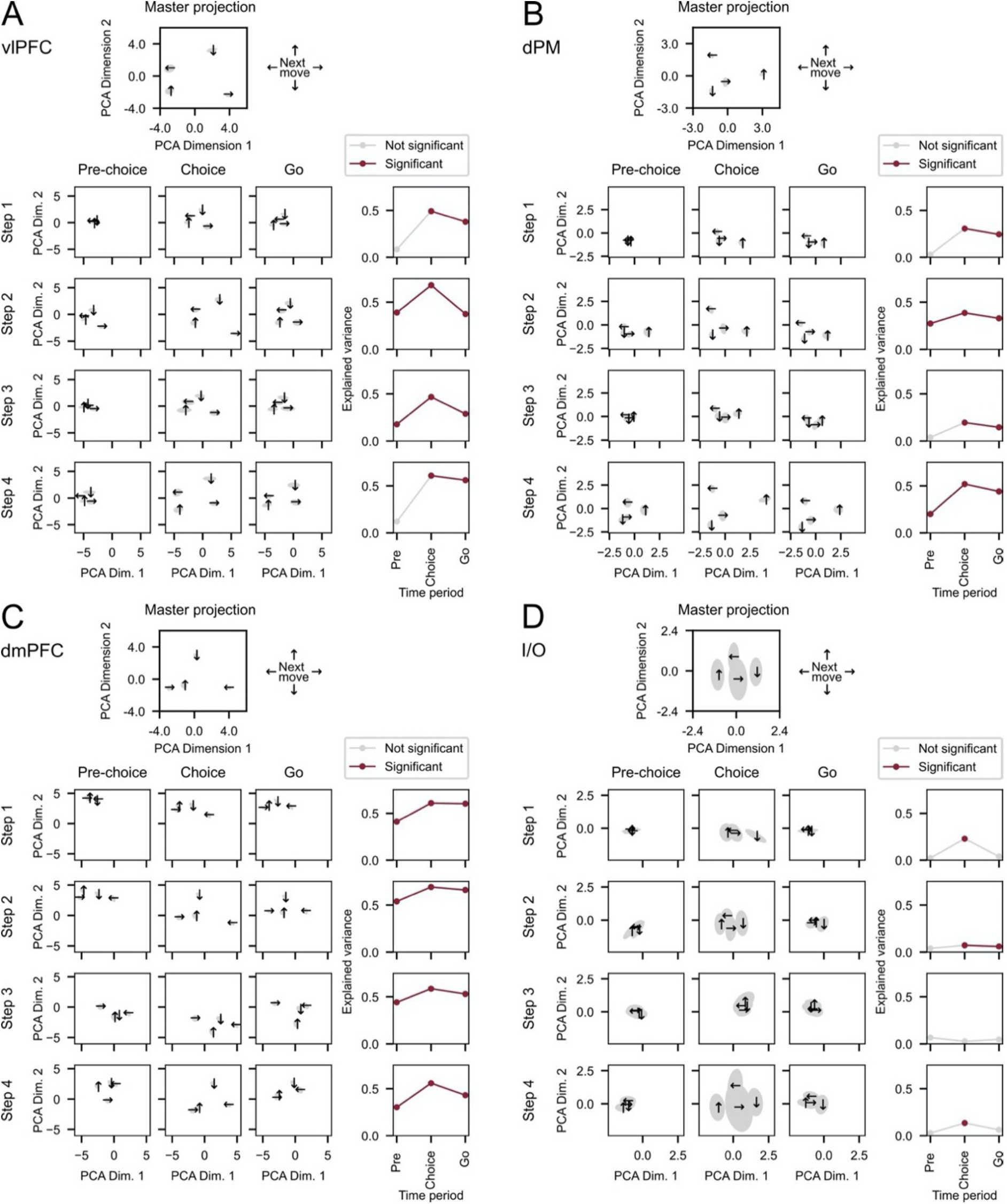
Move spaces for A - vlPFC, B - dPM, C – dmPFC, D – I/O. For each region, the top panel shows projections into a master space derived from data averaged over four goal locations for each move (choice periods from steps 2 (2-step problems only) and 4). Lower rows show projections into this master space for data from individual time windows in each step (left three panels), with proportion of explained variance in each window to the right (purple, *p* < .05, FDR corrected). In each row, data are sorted according to the move made on this step. Step 2 row takes data only from 2-step problems. Ellipses show confidence intervals derived by bootstrap resampling.

Figure 5 (lower rows in each panel) shows projections for each problem step, sorting data for each step according to the move made on this step. For step 2, again, we show data just for 2-step trials (for 4-step problems, see below). Apart from weaker move coding in I/O, the results show broadly similar dynamics across regions. At step 1, as expected, projections for the four moves separated rapidly after onset of the choice stimulus (Figure 5, compare pre-choice with choice). In dmPFC, small but significant variations within the move space, even prior to onset of the choice stimulus, suggest preparatory bias towards the two possible moves associated with the current goal. At later steps, move coding was robust even prior to choice stimulus onset, indicating preparation of the forthcoming move. For step 2, the results show anticipatory selection of the move required in a 2-step problem, i.e. the move that would reach the final goal. For steps 3 and 4, the required move was 100% predictable. At each step, move coding strengthened after onset of the choice stimulus, and remained up to the go period.

For a closer examination of neural dynamics, we focused on step 1, when onset of the choice stimulus strongly drove move selection, and the transition into step 2, when the selected move for step 1 was complete and the expected step 2 move was prepared. For this analysis we used 50 ms windows, covering the ranges -300 to +400 ms from choice 1 onset, -200 to +100 ms from choice 1 go signal, and -400 to +100 ms from choice 2 onset. For each window, we projected data into the move space, as defined above, and measured the distance of projected data from canonical positions for the four moves (Figure 5, top panel for each region, “master projection”). In Figure 6, we plot distance from the canonical position for the selected move 1, the prepared move 2, and for comparison, the two other moves (baseline, matching neither move 1 nor 2).

**Figure 6.**
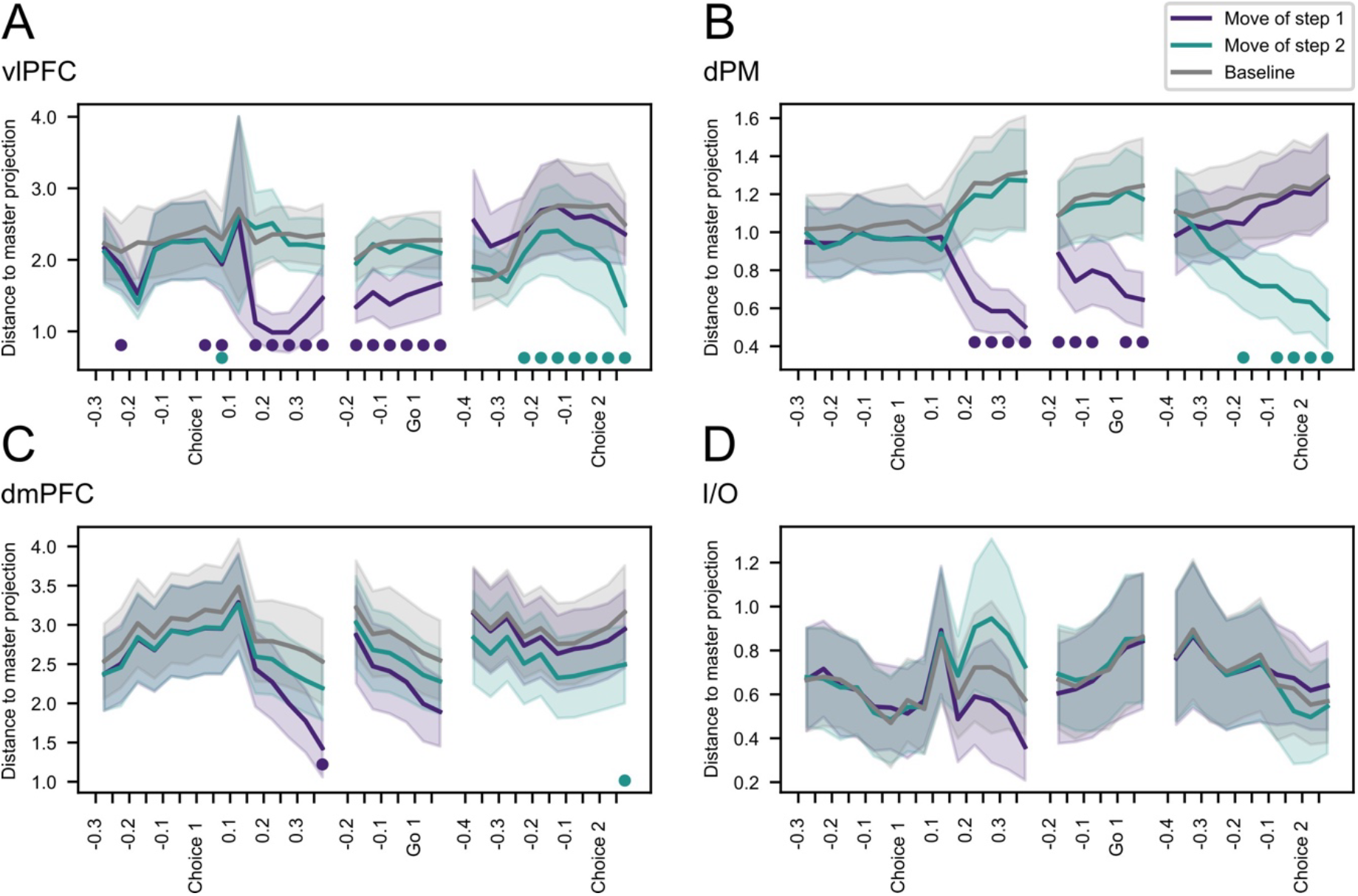
Distances of projected data from canonical positions in master move space, calculated for 50 ms windows spanning choices 1 and 2. A - vlPFC, B - dPM, C – dmPFC, D – I/O. Purple – distance of projected data from master position for the required move 1. Aqua – distance from master position for the required move 2. Grey – mean distance from master position for remaining two moves (baseline). Shading shows +/- 1 standard deviation over bootstrap samples. Bottom dots – difference from baseline significant at *p* < .05, FDR corrected.

Even prior to onset of the choice 1 stimulus, the data suggest weak bias towards both of the moves associated with the current goal (note that, for each goal, the step 2 move prepared following step 1 was also the alternative move that could itself have been cued for step 1; see Figure 1A). Though visible in vlPFC, dPM and dmPFC, these differences from baseline were small and mostly nonsignificant. Following onset of the choice stimulus, the representation shifted towards the selected move 1. This shift was seen first in vlPFC, followed by dPM and finally, more weakly, in dmPFC. Move 1 coding was sustained up to the time of the go signal, then followed by a rapid shift away from move 1 and towards the prepared move 2. In vlPFC, this shift towards move 2 was significant by 200-250 ms prior to choice 2, only 150-200 ms after completing the move 1 saccade (Figure 1A, right), with a slightly slower timecourse in dPM, and again, appeared only later and more weakly in dmPFC.

In 4-step problems, with onset of the choice 2 stimulus, the prepared move towards the goal had to be abandoned and a new move selected. Though move coding became disorganized (Supplementary Figure 3), by the time of the go signal, there were shifts away from the move initially prepared and towards the move selected.

Together, these results show that move coding in response to a visual input arose first in vlPFC, followed shortly afterwards by dPM. Move coding was seen also in dmPFC (Figure 4), but weak in I/O. Across regions, once a move was completed, there was rapid transition to preparation for the next.

### Orthogonality of spaces

To assess orthogonality of location and move spaces, we projected data for the four moves (window +200 to +400 ms) into the location space, and data for the four goals (window -200 to +100 ms) into the move space. In each case, the data projected were those used in the original definition of spaces, combining steps 2 and 4 to obtain a mean for each goal position, averaged over possible moves, and a mean for each move, averaged over possible goals. We asked what proportion of variance between the four moves was captured by projections in the location space, and vice versa. For vlPFC, dPM and I/O, spaces were close to orthogonal, with <8% of variance captured in these cross-projections (<5% for all cases except I/O, projection of moves into location space). For dmPFC, however, there was some overlap between spaces (22% of variance for move projected into location space, 28% for the reverse).

### Problem structure

Our final analyses concerned abstract problem structure separate from the specific content of any one problem. Again main analyses used combined data from the two animals. For an initial overview, we examined activity patterns for each successive period of a problem. For each of the periods previously defined (Figures 3, 5; three time windows for each step), and for each of the four goals, we calculated mean normalized spike rate for each neuron. For this analysis we also added two windows in the initial goal presentation period, 100-400 and 400-900 ms following onset of the goal stimulus. Vectors of spike rate across all recorded neurons were then correlated across two independent halves of the data (see Methods). The result (Supplementary Figure 4) was a matrix of correlations between activity patterns for each trial period x goal.

This overview of the data suggested three potential patterns of interest (Supplementary Figure 4). First, there was a strong diagonal, showing that patterns of activity changed substantially between each successive time window of problem solution. Second, there was a suggestion of hierarchical structure, with stronger correlations within than between successive steps, and between steps, stronger correlations for same versus different time windows. Importantly, these impressions remained even for correlations between problems with different goals, suggesting that they concerned abstract problem structure rather than specific problem content.

For formal analysis, we focused just on windows of steps 1 and 2 (2-step problems), for which there were most data. To focus on abstract problem structure, we considered only correlations between cases with different goals, current locations and moves (e.g. step 1, goal = upper left, current location = center, move = left correlated with step 2, goal = upper right, current location = above center, move = right; or step 2, goal = upper left, current location = left of center, move = up correlated with step 2, goal = upper right, current location = above center, move= right). Note that all correlations within step 1 were removed, since all step 1 cases had the same current location, leaving just retained step 1/step 2 and step2/step 2 pairs. For each retained pair, there was a 3 x 3 matrix of correlations (3 time windows for A x 3 time windows for B). Mean 3 x 3 matrices across retained pairs of each type are shown in Figure 7.

For all four regions, the impressions from the overall correlation matrix were confirmed. The strongest correlations were for same step, same window, confirming that, across the network, patterns of neural activity carry an abstract code of progress through the problem. Correlations for same step, different window, indicating a code of current step, and different step, same window, indicating a code of progress through the current step, were each stronger than the negative correlations for a baseline of different step, different window. For all regions, correlations for different step, different window were significantly lower than all others (*p* < .001 for all comparisons).

**Figure 7.**
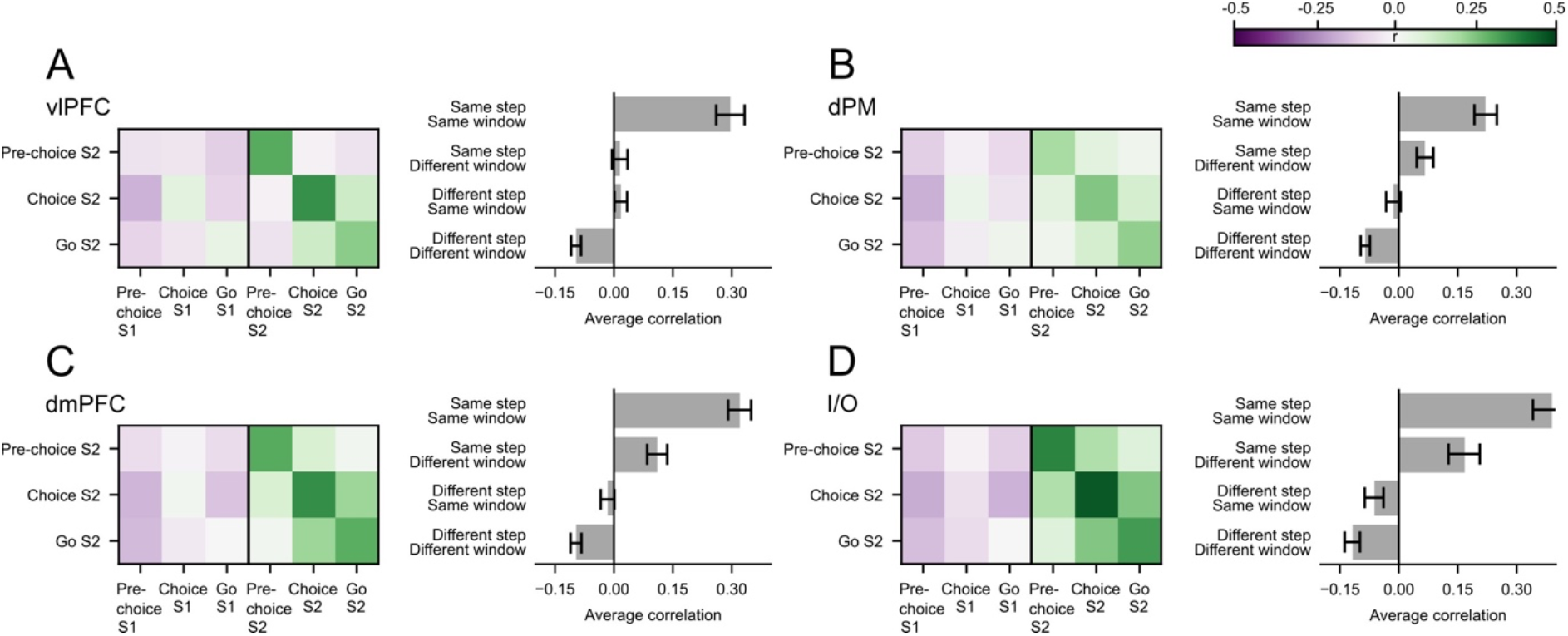
Correlation of cross-neuron activity patterns in A - vlPFC, B - dPM, C – dmPFC, D – I/O. For each region, left part shows mean correlations of each step 2 (S2) window with step 1 (S1) and step 2 (S2) windows. Data are means across all step 1/step 2 and step 2/step 2 pairs that have different goals, current positions and moves. Note that step 2/step 2 matrix is symmetrical (see Methods). Right shows mean correlations for same step same window, same step different window, different step same window, and different step different window. Whiskers show +/- 1 standard deviation over bootstrap samples.

Stronger correlations for same step, different window than for different step, different window might in part reflect simple temporal separation. To assess this, we compared correlations for step 2, pre-choice with step 1, go and step 2, go. Temporal separation for these cases was approximately matched (for monkey A, mean time between step 1 go and step 2 pre-choice 772 ms; mean time between step 2 pre-choice and step 2 go 900 ms; for animal 2, respectively 624 and 700 ms), but still, the correlation for step 2, pre-choice with step 2 go was significantly greater than the correlation for step 2, pre-choice with step 1, go (*p* < .02, .001, .005 and .001 respectively for vlPFC, dPM, dmPFC, I/O).

For vlPFC and dmPFC, separate data for the two animals are shown in Supplementary Figure 5, confirming similar findings in the two.

## Discussion

Cognitive control must rest on an internal task model, including specification of current state, goal state, and actions chosen to diminish the distance between the two. A salient feature is hierarchical structure, with a top goal controlling a series of subgoals. As monkeys plan a route through a maze, our data show all these properties in an extended frontal lobe network, putatively based on the extended MD network of human cognition. In four frontal lobe regions, there were strongly overlapping response properties, but at the same time, clear quantitative differences. The results suggest cognitive control by a network of partial frontal lobe specialists, together encoding the key features of a task model.

In vlPFC, we saw the strongest immediate coding of sensory input, including the target cue at trial onset, choice alternatives indicating the required first move, and an indication of current route type (2- or 4-step). vlPFC also strongly reflected current position in the maze, also associated with immediate sensory input. Orthogonal representational spaces in vlPFC coded current location and move, and compared with move spaces in other regions, vlPFC showed the most rapid change as a move was chosen. Previous findings have suggested vlPFC as an attentional hub ^27^, with strong coding of both spatial and nonspatial features of a current task ^28^ ^29^, and widespread cortical connectivity ^30^. Our data suggest a central role in the individual operations of each task stage – encoding a target, responding to sensory input and selecting an individual move.

In dmPFC, uniquely, there was a sustained signal of the final goal, beginning when this goal was first cued and lasting until problem end. Much previous research links dmPFC to reward processing (e.g. ^9^ ^31^ ^20^), based on extensive connectivity to reward systems ^32^. dmPFC has frequently been linked with energizing behavior ^33^ and a willingness to meet challenges ^34^ ^35^. An influential proposal links dmPFC to the tradeoff between prospective rewards and the effort needed to obtain them ^36^. In the maze task, a plausible possibility is that a sustained code of the goal location is linked to the reward obtained on reaching it. This sustained goal signal, we suggest, energises a series of steps or sub-goals along the path to its achievement (for a related proposal, see ^37^). Though a number of prior proposals have concerned the separate functions of dorsomedial and lateral frontal cortex (e.g. ^38^ ^39^), by separating goal coding from coding of individual task steps, we were able to show that one strong property of dmPFC is the stability of coding a higher-level goal.

In dPM, as in vlPFC, there was rapid move coding following onset of a choice stimulus, as expected for a motor region linked to primary motor cortex, and involved in both manual and saccadic movements ^40^. In I/O, there was weak coding of sensory input, locations and moves. Instead, there was strong response to input that overturned preparation for a short route to the goal, indicating a longer, more demanding path. Insula coding of progress towards a goal has been shown previously ^15^ ^19^. Plausibly, this signal may be related to the strong autonomic connectivity of the insula ^41^, and anticipatory arousal linked to goal proximity. Human fMRI data also show insula response to worse-than-predicted outcomes^42^.

Across regions, despite these partial specialisations, our data also showed substantial overlap of coding properties. Though vlPFC responded most rapidly to new sensory input, these rapid responses were followed by more slowly-developing activity across all regions, and for inputs indicating goal, move or route length. In dPM and I/O, there was coding of both current position in the maze, resembling vlPFC, and goal position, resembling dmPFC. Though motor signals emerged most rapidly in vlPFC and dPM, there was also a clear move space in dmPFC, and to some extent also in I/O. When moves were partially or wholly predictable, anticipatory move signals were seen across vlPFC, dPM and dmPFC. Strong interconnectivity between frontal regions ^23^ ^22^ provides a basis for substantial information sharing. Our data suggest that, in each region, there are often copies of information more strongly coded elsewhere. In each region, to a large extent, different aspects of the monkey’s problem were encoded in orthogonal state spaces, suggesting a medium for minimized interference between these distinct codes.

One common element across all four recording regions was a strong, hierarchical representation of abstract problem structure – the progress of steps from initial goal specification to final achievement. In each region, activity patterns were strongly correlated for the same time window in the same task step, even when goals, current positions and moves all differed. Patterns were more similar within than between task steps, suggesting a coarse, step-wise code of task structure. Between steps, patterns were more similar for the same than for different time windows, suggesting an abstract code of progress within each step, perhaps linked to the succession of required cognitive operations. Strong coding of the current point in progress from problem start to goal matches previous findings in both monkeys ^43^ and mice ^44^. Our data suggest that coding of abstract problem structure is a broad principle of prefrontal activity, perhaps providing a general scaffold onto which an individual problem solution can be attached ^45^.

Our results leave many questions open. Overlap in the response properties of different regions suggests the likelihood of communication, with each region influencing activity in the others. For example, the finding of early sensory responses in vlPFC might suggest that sensory information is distributed from this region to others. Current data, however, cannot test such hypotheses, as neural samples in a single session were generally not sufficient to show trial-wise transfer of information from one region to another. Our selection of regions as potential matches to human MD regions is tentative, especially for I/O where we have little direct evidence of correspondence. Though our data suggest partial specialisations between frontal regions – for example, strong coding of sensory input and current state in vlPFC, and stable goal coding in dmPFC – these specialisations call for examination in a wider range of multi-step tasks, including pursuit of nonspatial goals. The canonical human MD network includes regions outside the frontal lobe, conspicuously in lateral parietal cortex, and monkey data too show similar neural responses in lateral frontal and inferior parietal cortex ^46^ ^47^ ^48^ ^49^. Very likely, our data show only a part of a more distributed cortical and subcortical control network. We note too that results were heavily weighted towards recordings from one animal, though for vlPFC and dmPFC, data separated by animal showed similar trends.

Meanwhile, the data show how frontal components of this network encode essential aspects of cognitive control – goal, current state, component moves, and hierarchical problem structure. Resembling human imaging, these data suggest a network with partial specialisations, likely arising from distinct connectivity, but broad co-recruitment, likely reflecting information exchange. Evolving activity across this network of partial specialists builds the controlling task model for extended, goal-directed behavior.

## Methods

### Subjects

Subjects were two male rhesus monkeys (Macaca mulatta), each weighing 12 kg. Experiments were performed in accordance with the Animals (Scientific Procedures) Act 1986 of the UK; all procedures were licensed by a Home Office Project License obtained after review by Oxford University’s Animal Care and Ethical Review committee, and were in compliance with the guidelines of the European Community for the care and use of laboratory animals (EUVD, European Union directive 86/609/EEC).

### Task

In each session, the monkeys solved a series of problems defined in a 5 x 5 square maze, presented as a grid of lines on a computer screen (Figure 1A). Lines were white on a grey background. The 5 x 5 maze locations were defined by intersections between the lines of the grid. At each step of the solution, the current location was defined by the animal’s fixation on one of these intersections.

Moves consisted of a single step, i.e. a saccade from the current location to an immediately adjacent location. From a starting point in the central location, the task was to reach a specified goal location in a series of single steps. The goal was always located at one of the four corners of the inner 3 x 3 square of locations (Figure 1A, left). A correct trial consisted of taking the shortest route to the goal, considering the paths available at each choice. The types of trial were randomized between short (easy) consisting of only 2 saccades (2-step) before reaching the final goal, and long (difficult) with 4 saccades (4-step) (Figure 1A, left).

Timings are illustrated in Figure 1A (right). At the start of each problem, a dark green fixation point appeared at grid center. Fixation on this point was required within 3s. Once fixation was initiated, the dark green turned into bright green and after 100 ms a white diamond (cursor) joined the fixation point. After 200 ms a blue circle (target) appeared at the final goal location of the current trial and remained on the screen for 900 ms while the monkeys kept fixating in the center. Following the disappearance of this goal stimulus, there was a 300 ms delay, and then the four choice options of the first step were presented. Two dark green dots and two red dots appeared simultaneously at the four locations immediately adjacent to the central one (where the monkeys were still fixating). Red dots indicated steps unavailable for selection on this trial. Green dots indicated available choices, one of which was correct as it indicated a move towards the final goal. Following a delay period (randomly 300, 900, or 1500 ms for monkey A; 500 or 900 ms for monkey B), disappearance of the green fixation point served as a go signal. For monkey B, this go signal was accompanied by a small drop of reward (fruit smoothie). Following the go signal, the monkey had 500 ms (monkey A) or 350 ms (monkey B) to initiate a saccade to the correct choice location and 30 ms (monkey A) or 20 ms (monkey B) to complete it. Once the saccade was complete, if the choice was correct, the new fixation point changed to bright green and all the other old stimuli (center fixation point, white cursor, other choices) disappeared. While the monkey held fixation, the second step of the trial began. The cursor appeared after 100 ms and the second set of 4 choices appeared after a further 300 ms, presented for the same variable delay as before and followed by the go signal and saccade.

In 2-step trials, one of the choice 2 options lay at the final goal location, allowing the trial to be completed (Figure 1A, left, orange routes). In this case, after the correct saccade was made, the blue final target reappeared and a smoothie reward was delivered. On 4-step trials, however, monkeys needed to take a detour to the outer ring of the grid (Figure 1A, left, green routes). In this case, at step 2, the final target location was colored red, along with the previous fixation point at maze center. From the remaining two green choices, the monkey was required to choose the step to the outer ring. The third and fourth steps then followed the same structure as steps 1 and 2, though in each case, to enforce the correct route (Figure 1A, left, green routes), there was only one red dot (in the location just left) and two green. At the end of step 4, when one of the green choices lay at the final goal location, the trial ended as before with reappearance of the blue target and reward delivery.

On completion of each trial, all stimuli except the grid disappeared and there was an intertrial interval of 1800 ms. Throughout the problem, any error (wrong choice; fixation break; late response) led to problem termination, with a red error cue at screen center and no reward.

Task events were controlled by REX real-time data acquisition and laboratory control software (developed by the National Institutes of Health), with displays presented on a 44 cm LED screen placed 30 cm in front of the monkey’s eyes. The 5 x 5 maze measured 32 x 32 deg visual angle. Fixation point and red/green choice cues each measured 1.5 x 1.5 deg, while blue targets measured 3 x 3 deg. Eye position was sampled at 120 Hz using an infrared tracking system (ASL EYE-TRAC; Applied Science Laboratories www.asleyetracking.com), with each correct fixation required within a window of 3 x 3 deg. The eye tracker was re-calibrated at the start of each session.

### Surgery and recordings

For each animal, in two surgeries, a titanium head holder and recording chamber (Gray Matter Research) were sterotaxically implanted on the skull. Chambers were custom designed from an MRI based skull model of each monkey and placed over the lateral prefrontal cortex of the left hemisphere, giving access to dorsomedial and lateral frontal surfaces, along with anterior insula and adjacent orbitofrontal cortex. In a third procedure, a craniotomy was performed under each chamber to mount a 128-channel semichronic microdrive (LS-128-FL, Gray Matter Research). All surgical procedures were aseptic and carried out under general anaesthesia. At the end of the experiments, animals were deeply anesthetized with barbiturate and then perfused through the heart with heparinized saline followed by 10% formaldehyde in saline. The brains were removed for histology and recording locations confirmed.

Within the microdrive, each of the 128 single-contact tungsten electrodes was independently and manually movable in depth. Spacing between electrodes was 1.5 mm. The microdrive was interfaced with a multichannel data acquisition system (Cerebus System, Blackrock Microsystems). Raw extracellular signals were amplified, filtered (250 Hz to 5 kHz), and recorded in reference to the titanium head holder. Single action potentials were extracted and saved together with behavioral event codes in a file used for ofline sorting and analysis.

Initial spike sorting was performed online through the Blackrock interface before the start of each recording session. Sorting was then adjusted while the monkeys were performing the task and finalised ofline (Plexon Ofline sorting, version 4.7.1). In most cases, to ensure recording of new neurons, electrodes were advanced at least 62.5µ before every session. If this was not done, a Matlab-based algorithm was used to remove cases of repeat recording from the same neuron (https://uk.mathworks.com/matlabcentral/fileexchange/30113-tracking-neurons-over-multiple-days).

## Data analysis

Analysis scripts were written in Matlab (MathWorks) and Python.

To assess sensory information coding (Figure 2), for each cell, firing rates were examined by ANOVA with factors goal (goal presentation period), goal x horizontal/vertical choice available (choice 1 period) and goal x 2- vs 4-step (choice 2 period). From each ANOVA we calculated proportion of explained variance (PEV) for the factor of interest, and averaged this across all cells in each region. PEV was measured by the partial ω² index of effect size, calculated by the formula

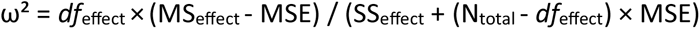

where *df*_effect_ is degrees of freedom for the factor of interest (target, choice, problem type), MS_effect_ is the mean square for the factor, SS_effect_ is the sum of squares for the factor, MSE is the mean square error, and N_total_ is the total number of observations (problems). Within each region, randomization tests were used to assess the significance of mean PEV values. For each neuron, labels for the factor of interest were randomized across problems, followed by re-calculation of mean PEV across cells. The PEV value for the true data was compared with the distribution obtained from 1000 repetitions of this randomization, and the *p* value determined by the proportion of randomized values greater than or equal to the true value. This procedure was repeated for each time window, and for each region, the criterion for significance was set at *p* < .05 corrected for multiple comparisons (time windows) by false discovery rate (FDR; ^50^). A similar procedure was used to compare PEV between regions, this time randomising just the region labels of each neuron.

To construct location and move spaces (Figures 3, 5), we used principal components analysis (PCA). For each neuron, missing data for a combination of location x move were filled in with the mean of other values. 95% confidence intervals (Figures 3, 5, ellipses) were constructed by bootstrap resampling of problems for each neuron. For each analysis window, we created 1000 bootstrapped projections, aligned by Procrustes transformation to the original projections.

To assess how much variance each space explained in successive problem periods (Figures 3, 5, rightmost panel in each row), we used a fully cross-validated approach. Two parallel data sets were created by randomly separating problems for each neuron into two groups, and PCA spaces were created for each half of the data. Data for each half were then projected onto the first two components of the PCA from the other half, and then reverse projected into the original data space. We then calculated the variance in the original data explained by the reconstructed data. Figures show the mean result from the two directions of cross-validation. To test significance, we repeated this process 1000 times while shufling condition labels in the projected data. The *p* value was given by the proportion of randomized values greater than or equal to the true value, FDR corrected for all comparisons for a given region.

To correlate activity patterns across problem periods (Supplementary Figure 4), problems for each neuron were split into two equally-sized sets. All firing rates for a given neuron were normalized (division by mean firing rate across the period from goal onset to end of step 2). Activity patterns for each region, and each half of the data, were then constructed as vectors of normalized firing rate across all neurons. Where a value for a given goal x time window was missing, it was filled in with the mean of that same time window for other goals. Similarity between vectors was measured by Pearson’s *r*.

To produce summary data (Figure 7), we took data from steps 1 and 2, with step 2 data just from 2-step problems. To focus on cases with different goals, current locations and moves, we began with a 48 x 48 correlation matrix, with rows and columns defined by 8 possible cases in step 1 (4 goals x 2 first moves), 8 possible cases in step 2 (4 goals x 2 current locations/next moves), and 3 time windows within each step. First, we removed all correlations between cases that shared goal, current location or move, including all step1/step 1 correlations (since in step 1, current location was always at maze center). Now there was no need to separate the data into independent halves, since the major diagonal (correlation of each case with itself) was removed. For each retained pair of cases (a step 1 case correlated with a step 2 case, or two different step 2 cases), there was a 3 x 3 matrix of correlations (3 time windows for case A x 3 time windows for case B). Mean correlations for step1/step2 and step 2/step 2 pairs (Figure 7) were calculated by *r* to *z* transformation. Note that, for step2/step 2, the 3 x 3 matrix of mean correlations is symmetrical since, for any pair of step 2 cases, it is arbitrary which is entered as row and which as column in the matrix; both assignments were accordingly used in calculating the average.

For analyses in Figures 4, 6 and 7, significance values were obtained by bootstrap resampling. For each of 1000 repetitions, we drew samples of neurons from each region, matched in size to the original data but sampled with replacement. The *p* value was given by the proportion of samples with a mean difference between conditions opposite to the majority, multiplied by 2 for a two-tailed test. The criterion for significance was set at *p* < .05, FDR corrected across time-points for each comparison in Figure 6.

## Data and code availability

Data and code are available at https://github.com/8erberg/Maze_review -- you will need this password to download the data: w+PQf-NpWd8n.GtJyZx+iW2

## Acknowledgments

This work was supported by Wellcome Trust Grant 101092/Z/13/Z and Medical Research Council UK Program MC_UU_00030/7. For the purpose of open access, the author has applied a Creative Commons Attribution (CC BY) licence to any Author Accepted Manuscript version arising from this submission.

## Supplementary figures

**Supplementary Figure 1.**
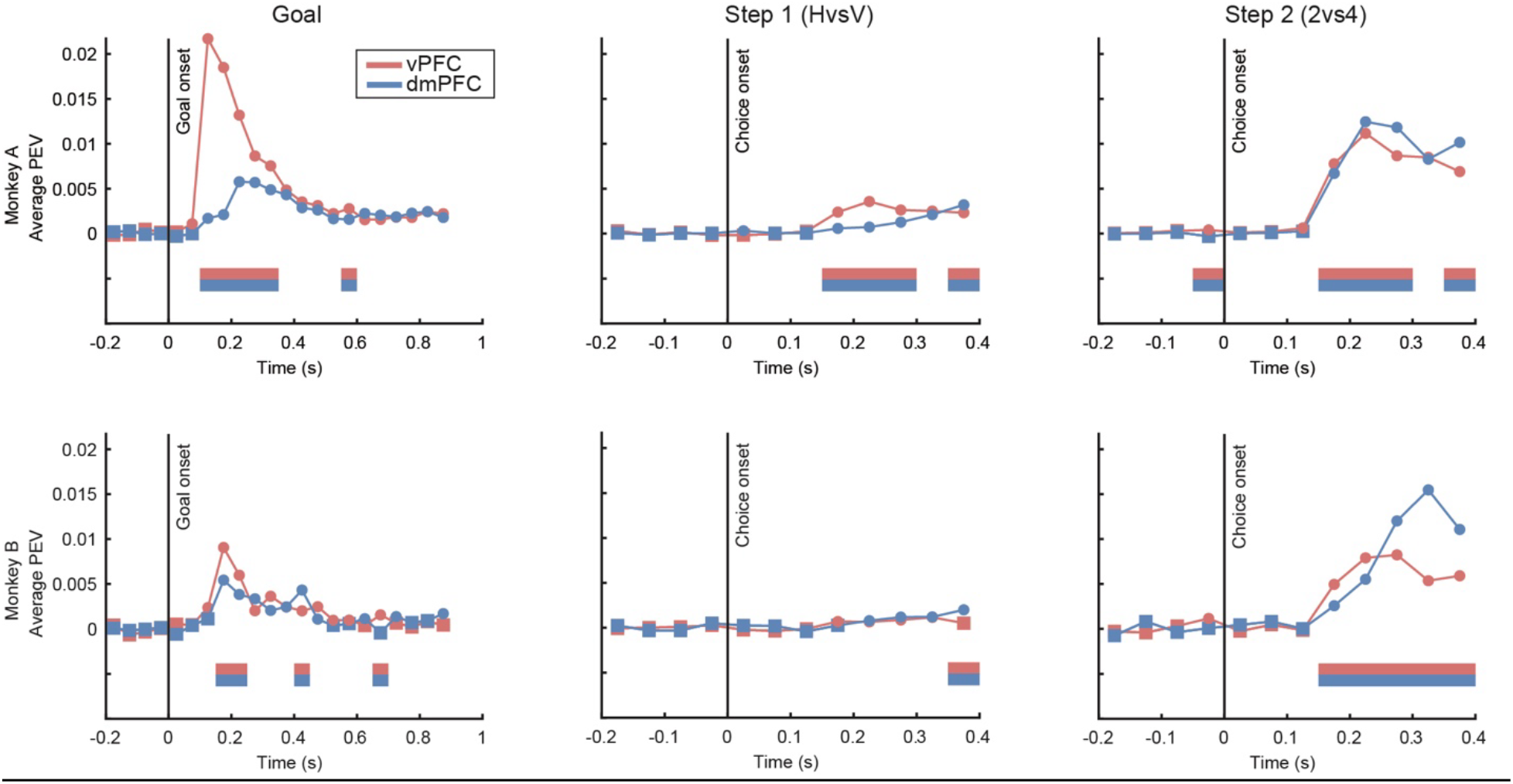
Mean proportion of explained variance (PEV) for goal (left, aligned at goal onset), horizontal (H) versus vertical (V) first move (middle, aligned at onset of choice 1 stimulus), and 2-versus 4-step problem (right, aligned at onset of choice 2 stimulus). Separate data fpr monkeys A/B. For each region, values of PEV significantly > 0 (*p* < .05, FDR corrected) are indicated by round dots, nonsignificant by square dots. Horizontal bars (bottom) show significant differences between each pair of regions.

**Supplementary Figure 2.**
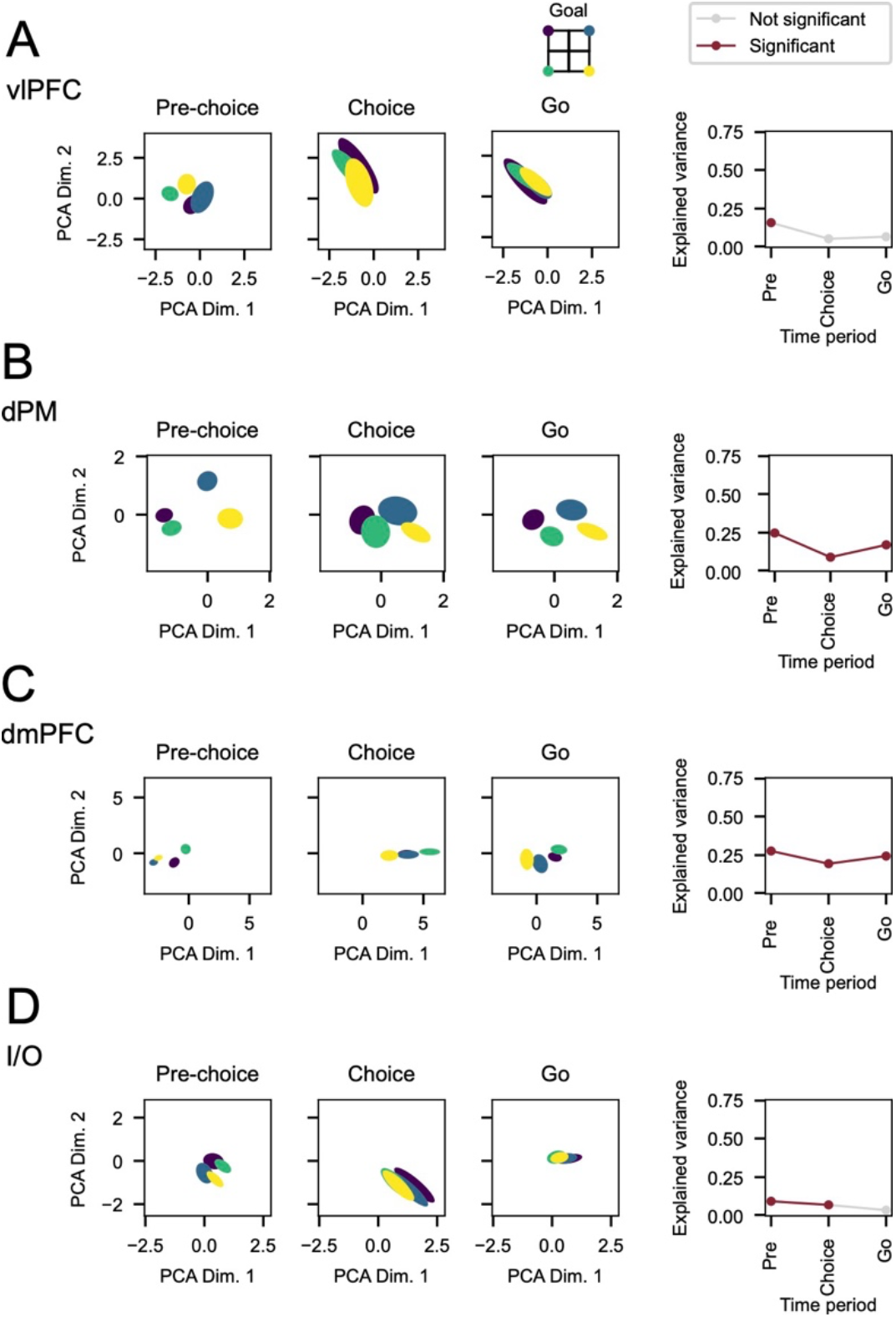
Projections of data from step 2, 4-step problems into location spaces. A - vlPFC, B - dPM, C – dmPFC, D – I/O. Conventions as fig. 3.

**Supplementary Figure 3.**
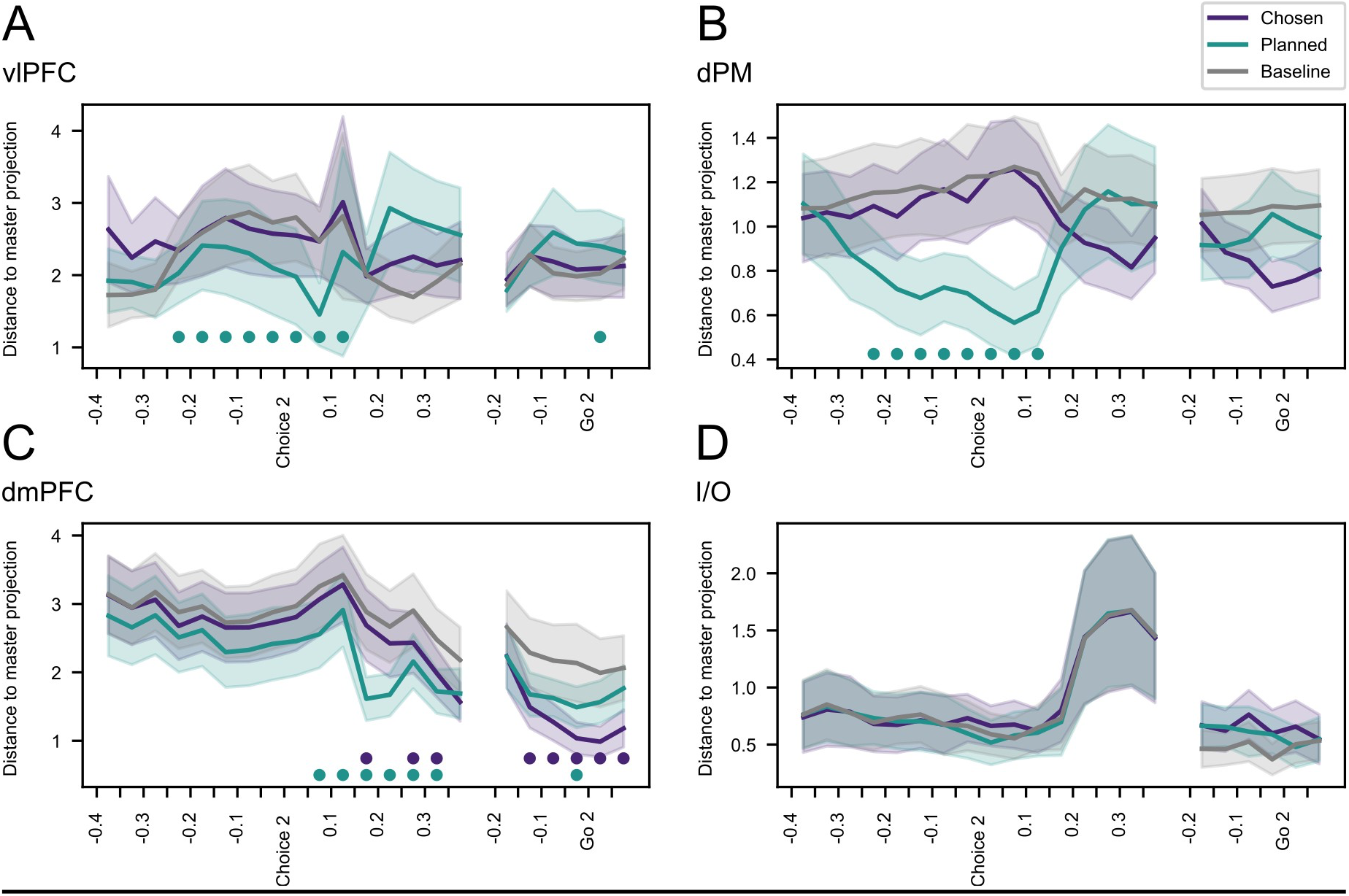
4-step problems: Distances of projected data from canonical positions in master move space. Data are shown for 50 ms windows spanning choice 2. A - vlPFC, B - dPM, C – dmPFC, D – I/O. Purple – distance of projected data from master position for the required 4-step move. Aqua – distance from master position for the planned 2-step move. Grey – mean distance from master position for remaining two moves (baseline). Shading shows +/- 1 standard deviation over bootstrap samples. Bottom dots – difference from baseline significant at *p* < .05, FDR corrected.

**Supplementary Figure 4.**
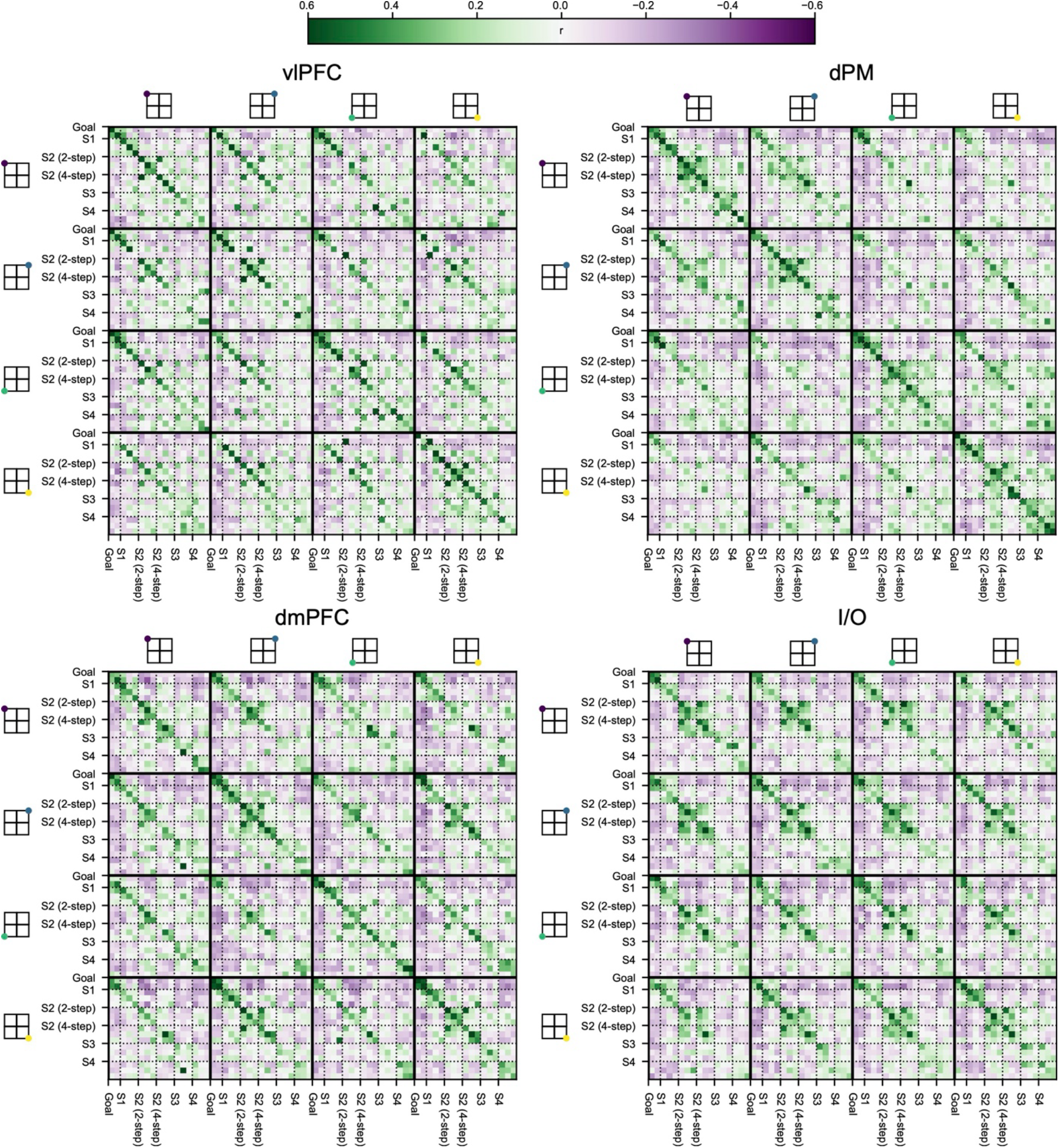
Correlation of cross-neuron activity patterns. A - vlPFC, B - dPM, C – dmPFC, D – I/O. From top to bottom, 17 time windows for each goal include 2 for goal presentation (100-400 and 400-900 ms post onset), and 3 each (pre-choice, choice, go) for step 1, step 2 2-step, step 2 4-step, step 3, step 4 (S = step).

**Supplementary Figure 5.**
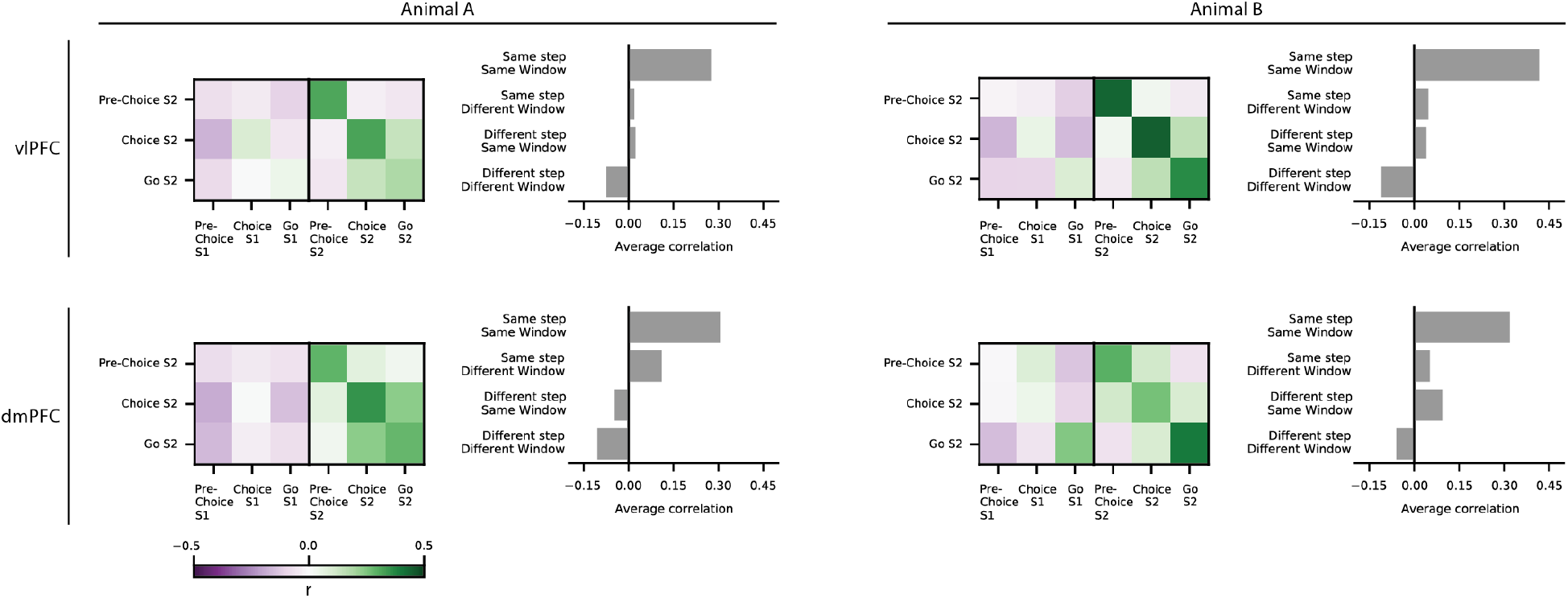
Correlation of cross-neuron activity patterns in A - vlPFC, B - dmPFC, separated by animal. For each region, left part shows mean correlations of each step 2 (S2) window with step 1 (S1) and step 2 (S2) windows. Data are means across all step 1/step 2 and step 2/step 2 pairs that have different goals, current positions and moves. Note that step 2/step 2 matrix is symmetrical (see Methods). Right shows mean correlations for same step same window, same step different window, different step same window, and different step different window.

